# Human-Relevant Lead Exposure During Differentiation Alters the Transcriptomic Profile of Human Induced Pluripotent Stem Cell-Derived Cardiomyocytes

**DOI:** 10.64898/2026.07.13.738273

**Authors:** Kimberley E. Sala-Hamrick, Anagha Tapaswi, Andre Monteiro Da Rocha, Justin A. Colacino, Laurie. K. Svoboda

**Author notes:** **Address for Correspondence:** Laurie Svoboda, PhD, M6232 SPH II, 1415 Washington Heights, Ann Arbor, MI, 48109.

## Abstract

Cardiovascular disease (CVD) etiology is strongly influenced by lead (Pb) exposure, but the underlying molecular and functional mechanisms are unclear, particularly during development. Using human induced pluripotent stem cell (iPSC) derived cardiomyocytes, we examined the effects of human-relevant Pb exposure during ventricular cardiomyocyte differentiation on transcription at several time points. We used an established protocol that temporally modulates Wnt signaling to differentiate iPSCs into contractile cardiomyocytes and exposed cells to 0.5 µM, 5 µM Pb, or control conditions during the first eight days of differentiation. Gene expression profiling on days 1, 2, 6, and 15 revealed significant Pb-induced changes in gene expression and dysregulation of pathways related to heart development and function, epigenetic machinery, and mitochondrial function throughout differentiation. Using BMDExpress3 modelling software, we calculated gene and biological pathway-specific best fit benchmark concentrations (BMCs) and found gene expression changes induced by Pb that were unique by day of differentiation but corresponded to a similar and human-relevant median benchmark concentration of 0.2 µM for all days assessed. Overall, our findings provide evidence that transient Pb exposure during cardiomyocyte differentiation causes transcriptional changes in human cardiomyocytes that persist even after cessation of exposure, underscoring the need for further investigation into how Pb exposure may impact heart development and function.

## Introduction

Lead (Pb) is a highly toxic, naturally occurring element that is strongly associated with cardiovascular disease (CVD), but effects of Pb exposure on heart development are incompletely understood.[1] Adverse cardiovascular effects of Pb have been reported in both prospective and cross-sectional epidemiological studies.[2] In 2019, Pb exposure contributed to an estimated 5.5 million adult deaths from cardiovascular disease worldwide (10% of global deaths that year).[3, 4] Pb exposure is associated with hypertension, arrhythmias, increased risk of mortality from myocardial infarction, increased incidence and mortality from coronary artery disease, and increased risk of peripheral arterial disease.[5–9]

The Developmental Origins of Health and Disease (DOHaD) hypothesis posits that environmental factors during critical windows of development can impact health across the life course. Notably, the DOHaD hypothesis originated in three studies that found poor nutrition at an early age could influence the risk of heart disease later in life, suggesting that the heart may be particularly susceptible to the effects of adverse developmental exposures.[10–12] Human observational studies strongly suggest that early exposure to Pb may influence long-term cardiovascular risk. Several investigations have identified associations between maternal blood Pb levels and increased risk of congenital heart defects in the fetus.[13–17] Further, prenatal Pb exposure has been linked to early life increases in blood pressure in children.[18] Rodent studies provide further evidence of Pb’s deleterious cardiovascular effects. We recently reported that developmental exposure to Pb has persistent impacts on the mouse transcriptome and epigenome that could contribute to later cardiovascular dysfunction.[19–21] Our findings are in keeping with work from others showing that early life Pb exposure in mice leads to changes in cardiac development and pathological responses in adulthood.[22] However, the mechanistic basis for Pb-induced developmental cardiotoxicity in humans remains poorly defined, including the critical windows of exposure and how Pb alters gene expression at important milestones of cardiomyocyte differentiation. Using human induced pluripotent stem cells (iPSCs), we sought to understand how Pb exposure during human cardiomyocyte differentiation would alter developmental gene expression and transcriptional pathways.

To this end, we exposed human iPSCs to human environmentally relevant concentrations of Pb (0.5 and 5 µM) or control conditions during the first 8 days of ventricular cardiomyocyte-directed differentiation and examined transcriptomic changes throughout differentiation (days 0-15). We chose this exposure window to recapitulate an *in-utero* cardiogenesis timeline comprising the developmental period during which pluripotent cells commit first to mesodermal fate and then cardiac progenitor lineages, prior to terminal cardiomyocyte differentiation, the onset of spontaneous contraction, and functional maturation of cardiomyocytes (days 8-30).[23, 24]

We conducted RNA-Seq in Pb exposed iPSC-derived cardiomyocytes from a female patient donor and found pathway alterations related to cardiac structure and function, development, epigenetics, energy metabolism, and mitochondria. We also used benchmark concentration modeling and identified gene and pathway-specific benchmark Pb concentrations at each time point during cardiomyocyte differentiation, finding human exposure relevant changes across all days examined. Overall, these studies provide insight into how Pb exposure may disrupt the transcriptome of the developing human heart.

## Methods

### Cell culture: iPSC maintenance

iPSC line PENN002i-442-1 from a 24-year-old Caucasian female reprogrammed from peripheral blood mononuclear cells was obtained from WiCell (WiCell, Madison, WI, USA). To promote adhesion and growth, iPSCs were plated in 24 well or 6 well Corning plates on surfaces coated with Corning Matrigel Basement Membrane Matrix that was prepared in DMEM:F12 HEPES (#07-200-80, #09-761-146 #356234 and #11330032; Thermo Fisher Scientific, Waltham, MA USA). iPSCs were maintained on this matrix in StemMACS iPS-Brew XF Media (#130-104-368, Miltenyi Biotec, Bergisch Gladbach, Germany) in feeder free conditions and incubated at 37°C in a 5% CO2 incubator. Media was changed daily according to the manufacturer’s instructions. Colonies of iPSCs that showed any signs of spontaneous differentiation were marked and removed via aspiration. Cells around 80% confluency were dissociated non-enzymatically with Versene (#15040066, Thermo Fisher Scientific, Waltham, MA USA) and seeded at low densities (1:24) for maintenance of iPSCs.

### Cell culture: iPSC differentiation

To perform differentiation into cardiomyocytes, iPSCs from above were passaged with Versene and seeded at higher densities into monolayers (passaged at 1:1 to 1:4 depending on starting day of differentiation). When cells reached 80-95% confluency (1-3 days), differentiation into ventricular cardiomyocytes began according to a published GiWi protocol with minor modifications (**Figure 1**).[24, 25] In the first 8 days of differentiation, cells were maintained on basal media (RPMI medium with HEPES CAT# 22400-89 Gibco, Grand Island, New York, USA with supplements ascorbic acid #A8960 and BSA #A9418 Sigma Aldrich, St. Louis, MO, USA) and switched over to B27 media on day 8 until purification. Modulation of Wnt/B-catenin signaling was utilized for differentiation as previously described (**Figure 1**).[24, 25] To direct cells into a mesoderm fate, iPSCs were incubated on day 0 in basal media with the GSK-3 inhibitor, CHIR99021 (#4423), which activates Wnt signaling, and after 2 days, cells were incubated with IWP4 (#5214), which inhibits Wnt signaling and drives cells towards a cardiac mesoderm, in basal media (Tocris Bioscience, Avonmouth, Bristol, United Kingdom; media: RPMI-1640 with HEPES, #22400089 Gibco, Grand Island, New York, USA, supplemented with 0.5% bovine serum albumin, #A8806, and 0.2% L-ascorbic acid, #A92902 Sigma Aldrich, St. Louis, MO, USA). Cells were then given basal media without the addition of small molecules on days 4 and 6 and switched over to RPMI media supplemented with B27 on day 8 with B27 media changes every other day after this until purification (RPMI # 11875-093 and B27 supplement #17504-044; Gibco, Grand Island, New York, USA).

**Fig. 1:**
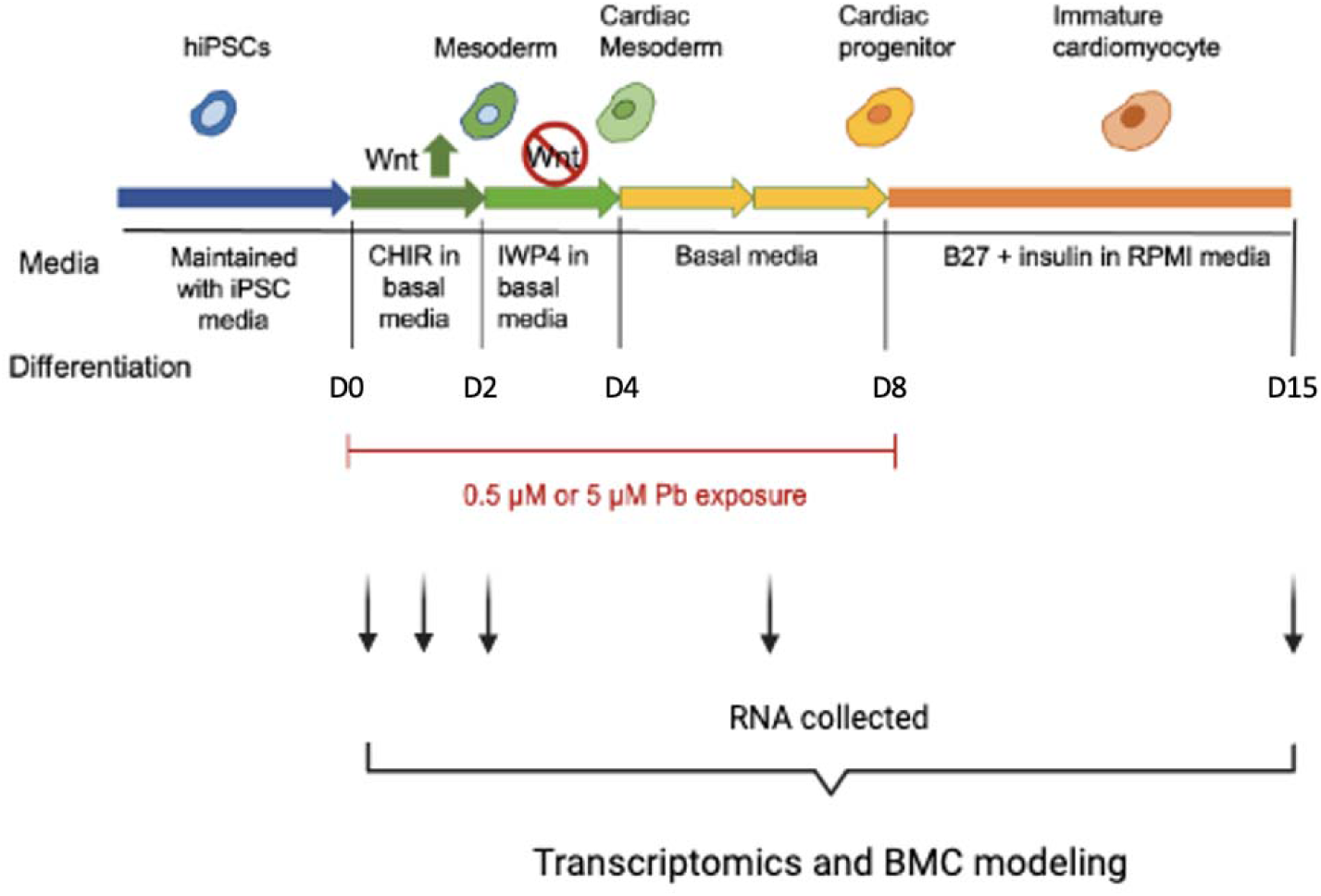
Protocol for differentiation of iPSCs into cardiomyocytes and experimental design. Ventricular cardiomyocyte differentiation protocol, with Pb exposure on the first 8 days of differentiation, RNA collected on days 0, 1, 2, 6, and 15.

### Cell culture: Pb exposure paradigm

As detailed in **Figure 1**, Lead (II) Acetate Trihydrate (CAS: 6080-56-4 ; #215902, Sigma-Aldrich, St. Louis, MO, USA) was diluted in tissue culture grade water and iPSCs undergoing differentiation were exposed for the first 8 days (days 0-8) at human exposure-relevant doses of 0.5 µM and of 5 µM. For 24 well plates (replicate 1) 3 wells were used for each exposure and combined for subsequent extractions or purification, while for 6 well plates, 1-2 wells were used for each exposure (replicates 2 and 3).

### RNA extractions

Cells were scraped off plates and pelleted before commencing RNA extractions. Cells were collected at days 0, 1, 2, 6, and 15 (**Figure 1**). RNA extraction was performed using the AllPrep DNA/RNA Universal Kit (#80224 Qiagen, Aarhus, Denmark) according to the manufacturer’s instructions with an additional DNAse incubation step (#79254, Qiagen, Aarhus, Denmark). Sample concentration and quality were assessed using a NanoDrop Microvolume Spectrophotometer (Thermo Fisher Scientific, Waltham, Massachusetts, USA).

### RNA sequencing: PlexWell cDNA preparation and quality control assessment

RNA sequencing was performed using the plate-based plexWell method (SeqWell), following a modified version of our previously established protocol.[26] RNA quantification was performed using the RiboGreen kit (Thermo Fisher, #R11490). For cDNA synthesis, 1 μL of RNA at 10 ng/μL was added to a 96-well plate (Dot Scientific, Cat. #951-PCR-B) and subjected to oligo annealing at 72°C for 10 minutes. Reverse transcription was then carried out at 50°C for 30 minutes, followed by an initial denaturation at 98°C for 3 minutes. Amplification proceeded through 12 PCR cycles of 98°C for 20 seconds, 67°C for 20 seconds, and 72°C for 6 minutes using a C1000 Thermal Cycler (Bio-Rad, Cat. #1851196). Samples were held at 4°C until purification. cDNA was purified using MAGwise Paramagnetic Beads (SeqWell, Cat. #MG10000), added in a 1:1 volume ratio with a 5-minute binding time. Samples were placed on a magnetic stand, the supernatant was removed, and beads were washed once with 80% ethanol (Spectrum, Cat. #E10258). Elution was performed in 20 μL of 10 mM Tris buffer (Thermo, Cat. #J22638-AE), and samples were stored at –20°C pending quantification.

cDNA concentration was measured using the Quant-iT PicoGreen dsDNA Assay Kit (Thermo, Cat. #P7589) following the manufacturer’s instructions. Samples and standards were plated in a 96-well format, and 100 μL of PicoGreen solution was added to each well. After incubation at room temperature in the dark, fluorescence was measured using the SpectraMax M5e and SoftMax Pro software v5.4. A global dilution factor calculator provided by plexWell was used to determine the appropriate cDNA input volume, targeting a minimum of 5 ng per sample.

### RNA sequencing: Library preparation using plexWell and sequencing

For library construction, 6 μL of cDNA at ∼1.7 ng/μL per sample was transferred to the sample barcode plate included in the plexWell LP384 Library Preparation Kit (SeqWell, Cat. #LP384X). Each sample was tagged with a unique i7 index using PCR conditions of 55°C for 15 minutes and 68°C for 10 minutes. After indexing, 18 μL of each barcoded sample was pooled, mixed with an equal volume of MAGwise beads, and incubated for 5 minutes to allow binding. Beads were pelleted on a magnetic stand and washed twice with 80% ethanol. Barcoded cDNA was eluted in 40 μL of 10 mM Tris.

Following pooling, sample concentration was re-quantified using the PicoGreen assay to ensure that the final concentration was within the recommended range (3.6–6 ng/μL). The pooled samples were then tagged with an i5 index (pool barcode) through a tagmentation reaction: 55°C for 15 minutes and 68°C for 10 minutes. After tagmentation, samples were purified using the previously described method and eluted in 24 μL of 10 mM Tris.

Final library amplification was conducted with the following thermocycling conditions: 72°C for 10 minutes, 95°C for 3 minutes, followed by 8 cycles of 98°C for 30 seconds, 64°C for 15 seconds, and 72°C for 30 seconds, and a final extension at 72°C for 3 minutes. Amplified libraries were diluted with 205 μL of 10 mM Tris, and 200 μL was purified using 0.8× volume of MAGwise beads. After magnetic separation and two ethanol washes, libraries were eluted in 30 μL of 10 mM Tris and stored at –20°C for short-term use.

Library quality was assessed using the Agilent Bioanalyzer with the High Sensitivity DNA 5000 Kit at the University of Michigan (UM) Advanced Genomics Core. Libraries were sequenced on the Illumina NovaSeq 6000 S4 platform with 151-bp paired-end reads, targeting an average depth of >30 million reads per sample.

### RNA sequencing: Data processing and analysis

Raw reads were demultiplexed using i7 and i5 barcodes. Sequencing data were transferred to the University of Michigan Great Lakes High-Performance Computing Cluster for analysis. Quality control was performed using FastQC and MultiQC.[27, 28] Reads were aligned to the GRCh38 human genome reference (splice-aware build) using STAR.[29] Gene-level counts per million (CPM) were calculated with featureCounts, excluding multi-mapped and multi-overlapping reads.[30]

### Computational analysis: DEG and pathway analysis

The R package DESeq2 was used to conduct differential gene expression analysis comparing control to Pb exposed samples individually for each day (0, 1, 2, 6, and 15) in female derived iPSCs undergoing differentiation into cardiomyocytes. Separate DESeq analyses were performed for each day, and replicate (batch) and Pb exposure were included in the model design. DEG analysis examined genes with adjusted p-values of less than 0.05. Principal component analysis (PCA) plots were made using the plotPCA in DESeq2. Gene annotation was performed using biomaRt.

To assess for pathway enrichment, the LRPath function was utilized as previously described.[31] This function employs logistic regression using the p-value and fold change for each gene (generated from DESeq2) to test for enriched or depleted biological categories, and GOBP (biological processes), GOMF (molecular function) and GOCC (cellular compartment) were chosen as the MsigDB gene sets for this analysis.[31] Significant pathways were selected with and FDR cutoff of 0.05. To select for relevant pathways, terms were selected and sorted into 5 categories: cardiac (heart|cardiac|ventric|atrium|muscle|atrial|aorta), mitochondria (mitochondri|respiratory|electron transport|oxidative), metabolism (ATP|glycolysis|fatty acid|glucose|metabolism), development (development|differentiation), and epigenetics (epigenetic|chromatin|histone|methylation|acetylation). The top three terms for each category per day/exposure were selected and plotted in a dot plot and the significance of all terms per category was plotted in a heat map using ggplot2.

### Benchmark concentration analysis

Benchmark concentration (BMC) analysis was performed as previously described for RNA sequencing data comparing control to 0.5 and 5 µM Pb on days 1, 2, 6, and 15 of differentiation.[26] Gene-specific benchmark concentration and best fit benchmark concentration models were identified using BMDExpress version 3.2, a free software package developed by the National Institute of Environmental Health Sciences (NIEHS) and available for download (https://github.com/auerbachs/BMDExpress-3/releases). Best practices for dose-response modeling for each chemical were conducted according BMDExpress online documentation (https://bmdexpress-2.readthedocs.io/en/feature-readthedocs/basic_workflow/). Normalized counts per million reads were imported into BMDExpress and prefiltered using one-way analysis of variance (ANOVA) for significance at a -value of <0.05 to identify genes showing significant increasing or decreasing concentration responses. To determine concentration– response relationships, the filtered data were modeled in BMDExpress with power, linear, polynomial (2°, 3°, 4°), and exponential (3 and 5) models and the best fit model was chosen. Best fit models were chosen for each concentration–response relationship per gene using nested chi square to select for the best model followed by lowest Akaike information criterion (AIC). The benchmark response (BMR) was set to 1 standard deviation relative to control response, maximum iterations of the model was set to 250, a confidence level of 0.95 was used, and power was restricted to ≥1 for the power model. Directionality in response (upregulated/downregulated) was derived from the “Best adverseDirection” of the winning model for a given gene. Benchmark concentrations with values above the highest tested concentration (5μM) were excluded from individual gene and functional classification analyses. We conducted functional classifications analysis for the three Gene Ontology sets (Biological Process, Molecular Function, and Cellular Compartment). Pathway analysis results were filtered at a significance at a Fisher’s exact two-tailed -value of <0.05 for BMDExpress testing. We selected for key words as described above for these pathways, and results were visualized by dot plot using ggplot2. All original expression data, ANOVA filtered results, BMC results, and subsequent BMDExpress analysis for all chemicals are included in a .bm2 file available as supplemental information.

### BMC and NHANES comparison

To compare transcriptomic BMC data to population level data, blood Pb concentrations were obtained from the most recent National Health and Nutrition Examination Survey (NHANES; 2021-2023) laboratory data. Participants with non-missing blood Pb measurements and complete survey design variables were included in the analysis and these values were converted from µg/dL into µM based on the molecular weight of Pb. The survey package in R was used to weigh the data according to survey design, including primary sampling units (SDMVPSU), strata (SDMVSTRA), and the appropriate two-year sample weight (WTPH2YR). Weighted blood Pb percentiles were then estimated using svyquantile. For the boxplots, BMC analyses were plotted alongside this NHANES data to contextualize the BMC modelling results according to population level data. For the first four boxplots, ggplot2 was used to plot days 1, 2, 6, and 15 of differentiation with the interquartile range (IQR) as the box and the whiskers representative of 1.5 x IQR range. For the NHANES boxplot, IQR was plotted as the box and the 1^st^ and 99^th^ percentiles used as the lower and upper whiskers, respectively.

## Results

### PCA plots show clustering by day of differentiation and replicate but not by Pb dose

To examine gene and pathway changes, RNA sequencing was conducted for iPSCs undergoing differentiation into ventricular cardiomyocytes on days 0, 1, 2, 6 and 15 of differentiation (n=3 independent experiments). Principal component analysis (PCA) plots were made across all days of differentiation and all doses, and showed clustering by day (**Figure 2**); (PCA plots for replicate and Pb concentration shown in **Figure S1**).

**Fig. 2:**
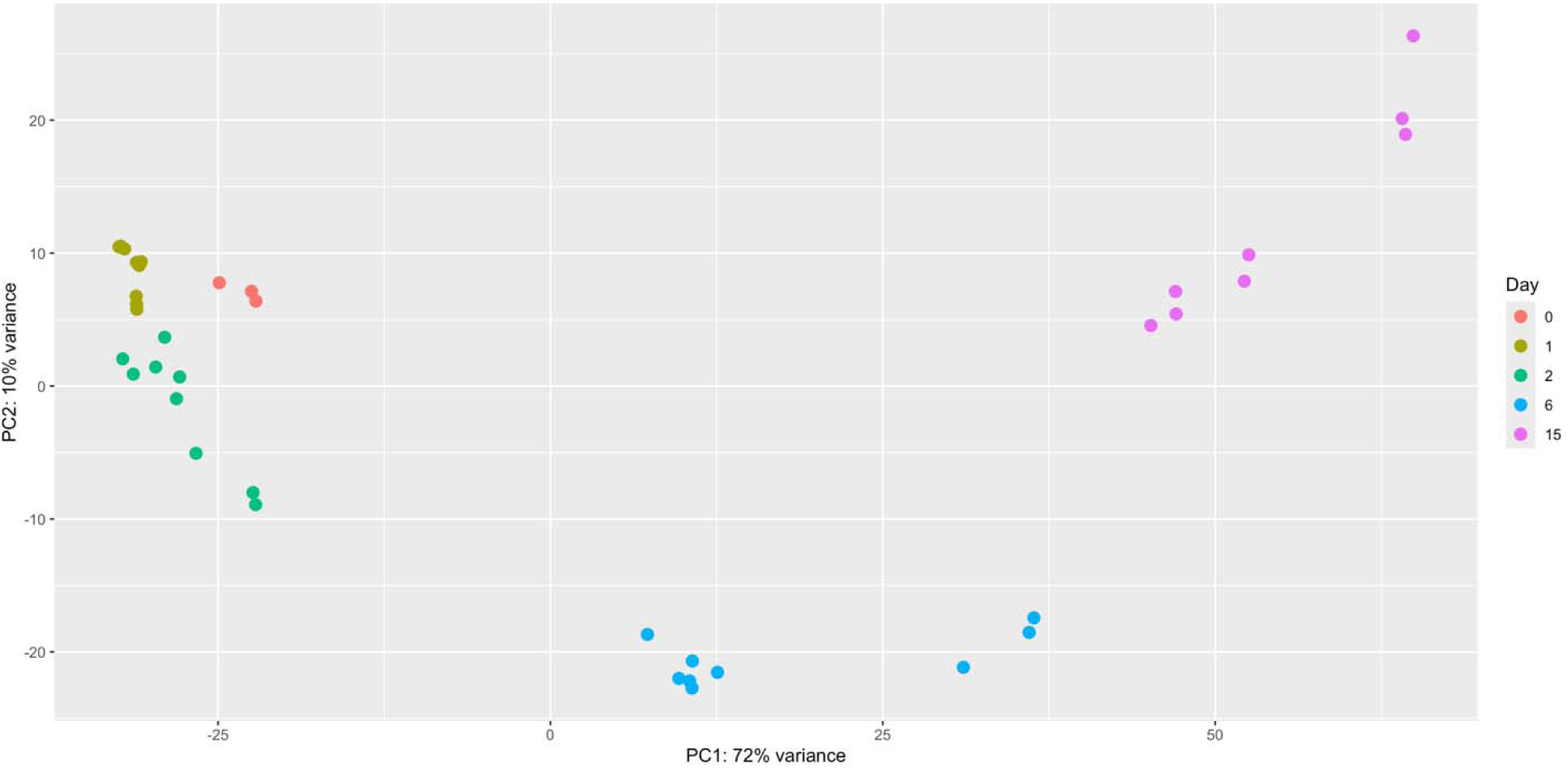
PCA plot reveals clustering by day of differentiation. Principal components analysis of RNA sequencing data of female iPSCs undergoing differentiation into cardiomyocytes, with points labeled by day of culture (0, 1, 2, 6, 15) (n=3 independent experiments per dose).

### Pb induces transcriptional changes throughout differentiation

We tested for differentially expressed genes separately for each day, comparing Pb exposure to control. We found a dynamic set of differentially expressed genes (padj > 0.05) that were unique to each day and Pb dose (full unfiltered gene expression results in **Table S1**). Interestingly, 5 µM Pb showed the greatest number of differentially expressed genes, with day 6 showing 77 genes up or downregulated compared to control (**Table 1**). **Figure S2** shows overlap in DEGs between all days and doses examined. Notably, the two DEGs overlapping between days 2 and 6 (5 µM) are *CCKBR* and *FOXA2*.

**Table 1:** Differentially expressed genes by Pb exposure in female iPSCs undergoing differentiation into ventricular cardiomyocytes. Differentially expressed genes identified using DESeq2 with a padj cutoff of 0.05 for 4 time points during differentiation (days 1, 2, 6, and 15) and 2 doses of Pb (0.5 and 5 µM) for cells exposed during the first 8 days of differentiation. Overlap between 0.5 and 5 µM doses within the same day are shown in the last row.

|  | D1_0.5 | D1_5 | D2_0.5 | D2_5 | D6_0.5 | D6_5 | D15_0.5 | D15_5 |
| --- | --- | --- | --- | --- | --- | --- | --- | --- |
| **Total** | 16 | 23 | 2 | 19 | 0 | 77 | 0 | 6 |
| **Up** | 8 | 11 | 0 | 1 | 0 | 7 | 0 | 0 |
| **Down** | 8 | 12 | 2 | 18 | 0 | 70 | 0 | 6 |
| **Overlap between**<br>**0.5 and 5μM** | 8 |  | 0 |  | 0 |  | 0 |  |

### Gene ontology analysis reveals Pb associated pathway perturbations that are dynamic throughout differentiation

We next wanted to investigate the transcriptional pathways impacted by Pb exposure during differentiation of iPSC into cardiomyocytes. Using LRPath, we tested for gene set enrichment in GOBP (biological processes), GOMF (molecular functions) and GOCC (cellular compartments) in our data and selected significantly altered pathways (FDR <0.05; all results shown in **Table S2**).[31] **Table 2** displays the top 30 pathways ranked by enrichment FDR across all doses and days for all three analyses, sorted by day/dose and including repeats in pathways between days/doses. 18 pathways out of 30 pertain to ribosomal function, while 8 out of 30 pertain to mitochondrial pathways or function. Relevant cardiac, developmental, epigenetic, metabolic, and mitochondrial terms were further selected and the top three terms for each day/dose are shown in **Figure 3**. Interestingly, pathways related to cardiac differentiation were found to be changed by Pb exposure in later stages of differentiation (striated muscle cell tissue development for 5 µM on day 15, stem cell differentiation for 5 µM on day 6, and cardiomyocyte differentiation for 0.5 µM on day 6; **Figure 3c** and **d**). Heat maps of mean p-values for each pathway reveal dynamic pathway regulation throughout differentiation, with highly significant changes in mitochondrial pathways on days 1, 6 and 15, but not on day 2 (**Figure 3e**).

**Fig. 3:**
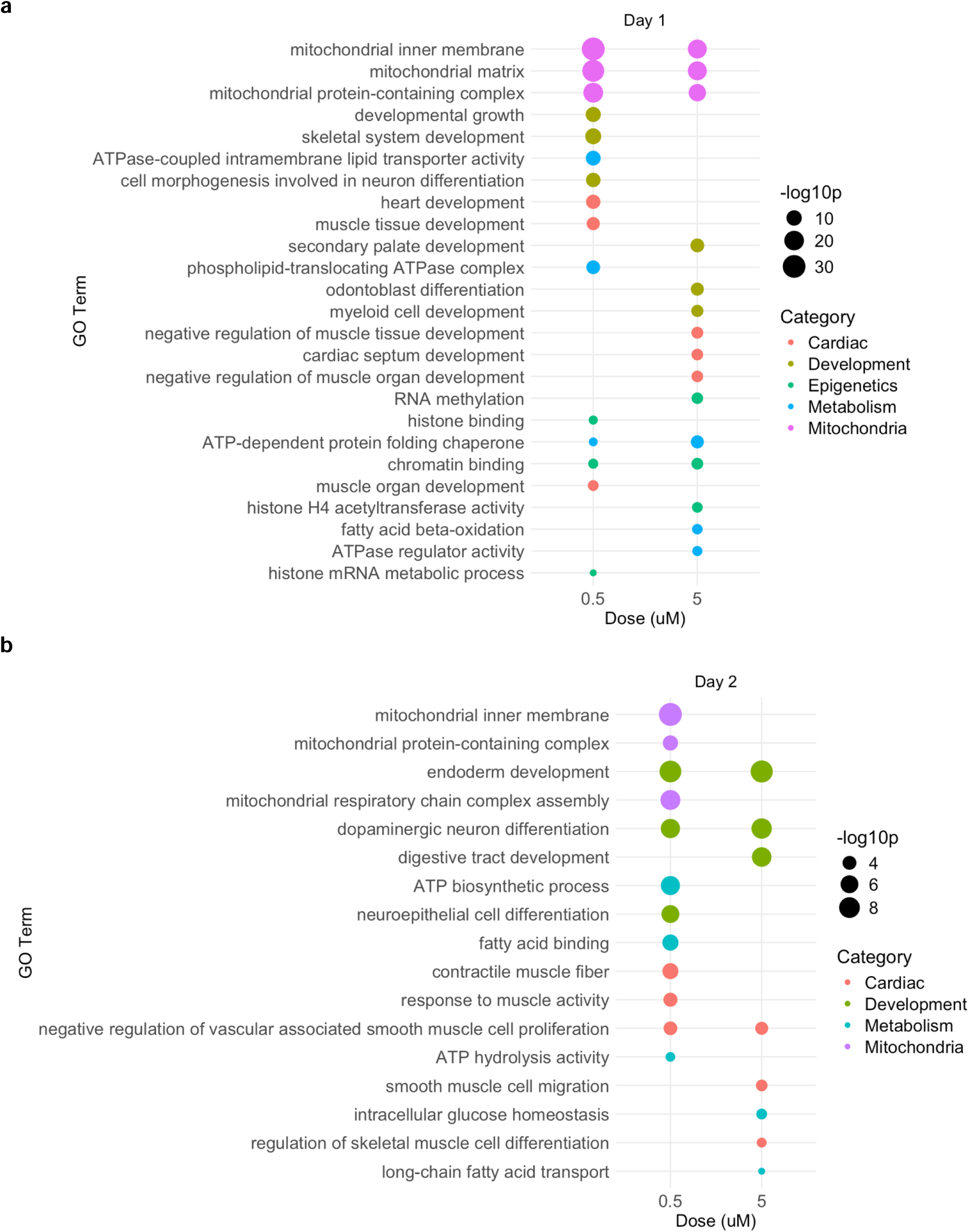
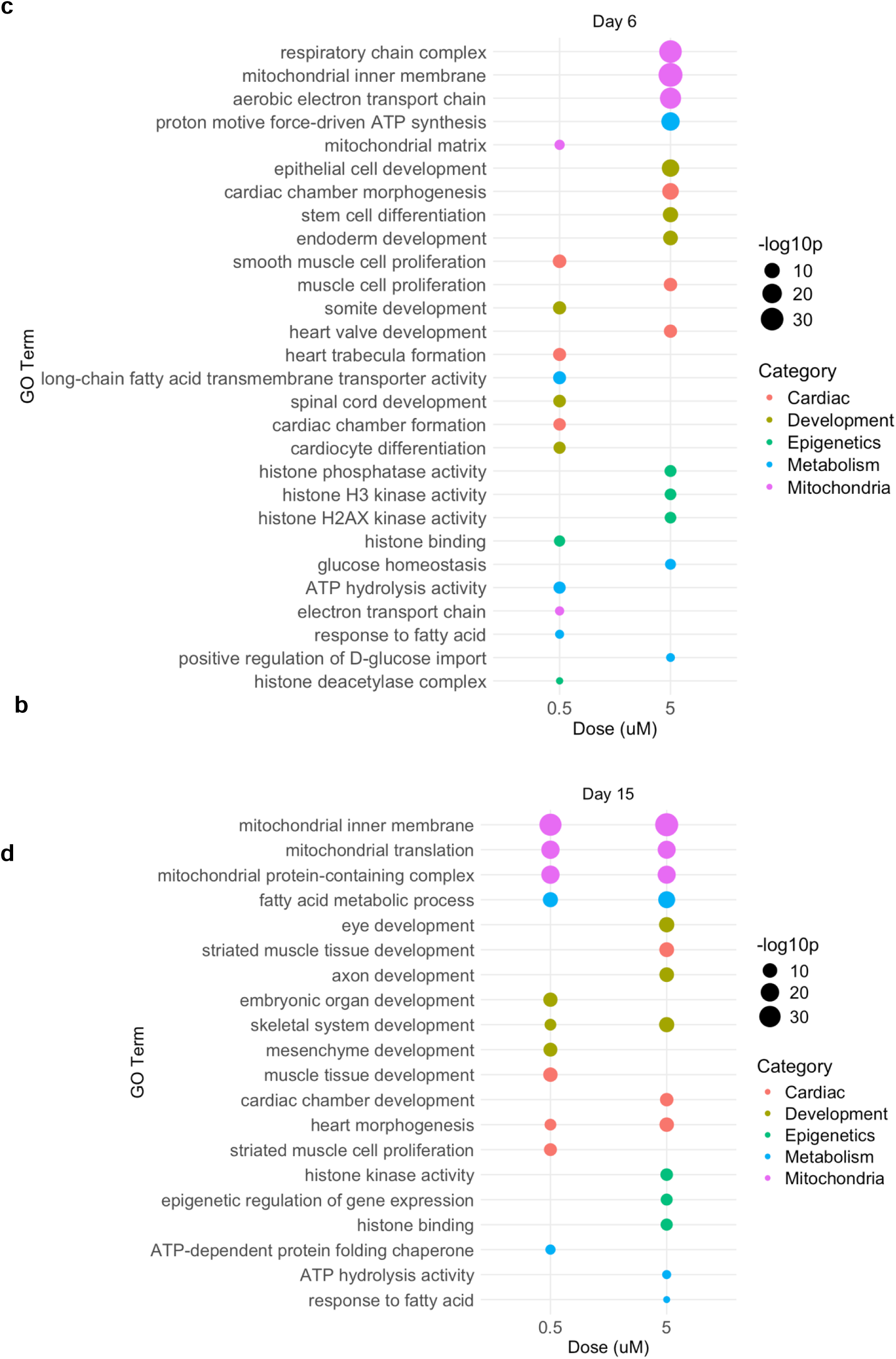
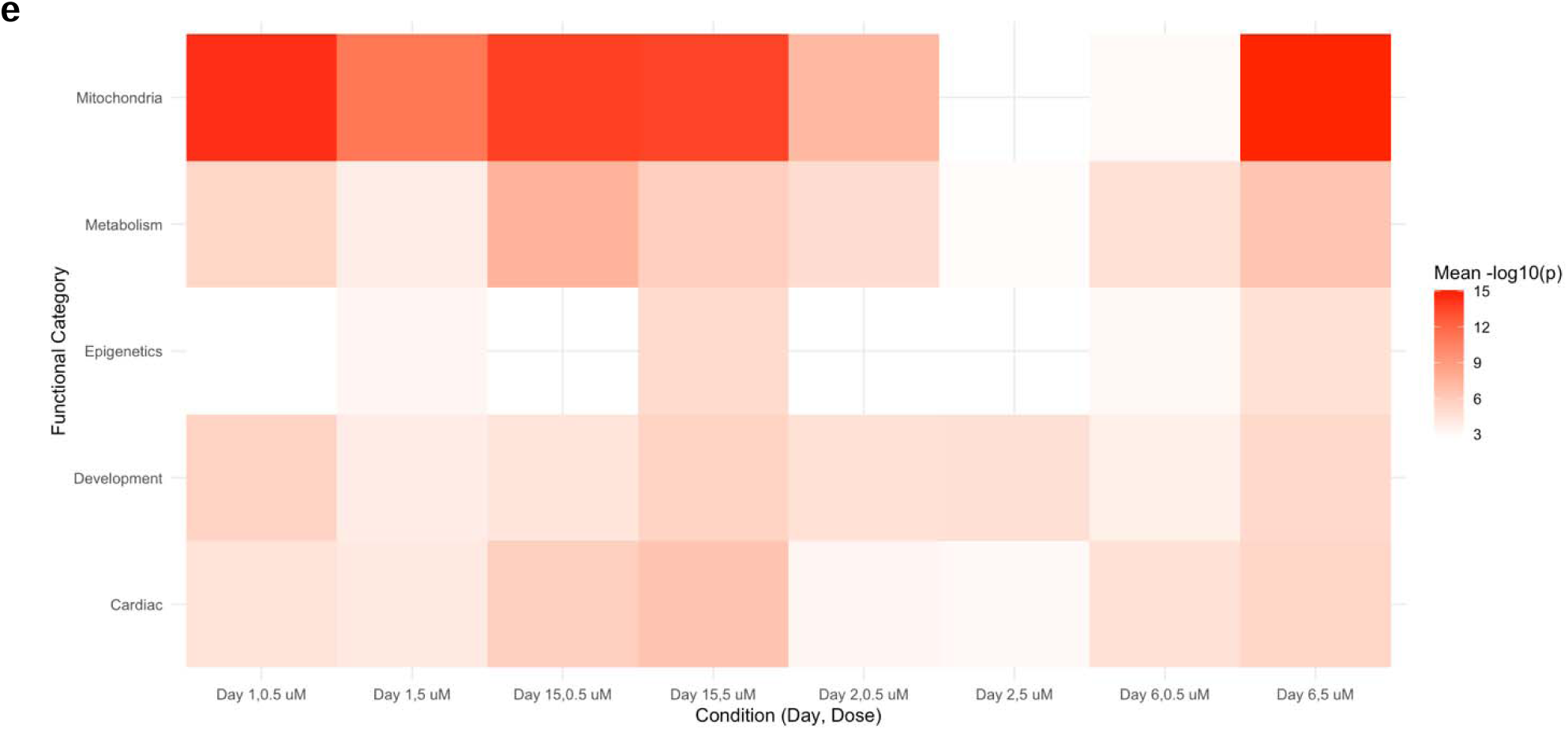
Pathways related to important cardiac processes were altered by Pb exposure during differentiation of iPSCs into ventricular cardiomyocytes. LRPath results for GOBP, GOCC, and GOMF divided into the top 3 terms for cardiac, development, epigenetics, metabolism, and mitochondria relevant categories for days 1 (a), 2 (b), 6 (c), and 15 (d) of ventricular cardiomyocyte differentiation with two doses of Pb (0.5 and 5 µM). The size of dot in each dot plot represents -log10(p-value) and the color corresponds to each category. Heat map of mean(-log10(p-value)) for each category across all days and doses is shown in (e).

**Table 2:** Unbiased top 30 GOBP, GOMF, and GOCC pathway alterations across days and doses. The top 30 pathways ranked by FDR for the LRPath results for GOBP, GOCC, and GOMF for days 1, 2, 6, and 15 of ventricular differentiation with two doses of Pb (0.5 and 5 µM), sorted in the table by day/dose.

| Description | Day | Dose (μM) | FDR |
| --- | --- | --- | --- |
| ribosomal subunit | 1 | 0.5 | 3.81E-59 |
| structural constituent of ribosome | 1 | 0.5 | 6.67E-57 |
| translation | 1 | 0.5 | 1.59E-54 |
| ribonucleoprotein complex biogenesis | 1 | 0.5 | 7.94E-52 |
| mitochondrial inner membrane | 1 | 0.5 | 1.28E-29 |
| rRNA metabolic process | 1 | 0.5 | 1.75E-29 |
| mitochondrial matrix | 1 | 0.5 | 3.28E-26 |
| spliceosomal complex | 1 | 0.5 | 3.62E-24 |
| RNA splicing, via transesterification reactions with bulged adenosine as nucleophile | 1 | 0.5 | 7.56E-24 |
| preribosome | 1 | 0.5 | 4.06E-20 |
| ribosomal subunit | 1 | 5 | 1.20E-33 |
| structural constituent of ribosome | 1 | 5 | 2.86E-28 |
| translation | 1 | 5 | 4.76E-24 |
| ribonucleoprotein complex biogenesis | 1 | 5 | 5.05E-22 |
| structural constituent of ribosome | 6 | 5 | 2.05E-49 |
| ribosomal subunit | 6 | 5 | 1.19E-47 |
| cytosolic ribosome | 6 | 5 | 2.10E-42 |
| mitochondrial inner membrane | 6 | 5 | 2.35E-33 |
| cytoplasmic translation | 6 | 5 | 1.64E-29 |
| respiratory chain complex | 6 | 5 | 1.58E-28 |
| cell-cell junction | 6 | 5 | 1.31E-24 |
| aerobic electron transport chain | 6 | 5 | 1.98E-23 |
| ATP synthesis coupled electron transport | 6 | 5 | 2.33E-23 |
| angiogenesis | 6 | 5 | 3.24E-23 |
| ribosomal subunit | 15 | 0.5 | 2.98E-37 |
| structural constituent of ribosome | 15 | 0.5 | 9.18E-32 |
| mitochondrial inner membrane | 15 | 0.5 | 3.40E-31 |
| mitochondrial inner membrane | 15 | 5 | 3.96E-35 |
| ribosomal subunit | 15 | 5 | 3.64E-31 |
| structural constituent of ribosome | 15 | 5 | 4.33E-25 |

### Benchmark concentration modeling reveals unique gene and pathway expression patterns throughout differentiation at similar benchmark concentrations

To understand the concentration of Pb that induces changes in gene or pathway expression during differentiation into ventricular cardiomyocytes, we performed benchmark concentration (BMC) modeling. This approach fits concentration–response models to the gene expression data comparing control vs. 2 concentrations of Pb. It then determines a BMC at which there is a change in gene expression equal to 1 standard deviation above or below the mean expression level in the control group. Genes with modeled responses meeting this benchmark response (BMR) and showing adequate model fits were retained for downstream analysis. BMC modeling using BMDExpress3.2 revealed that of all genes meeting these criteria, there were similar median BMCs on each day of differentiation of around 0.2 µM (**Figure 4**). The 5^th^ boxplot in **Figure 4** shows the weighted NHANES blood Pb distribution in the United States 2021-2023. Further, each day showed differences in the total number of genes dysregulated and those that were upregulated or downregulated (**Table 3**), with no overlap between days in the genes showing a BMC relationship (**Figure S3)**.

**Fig. 4:**
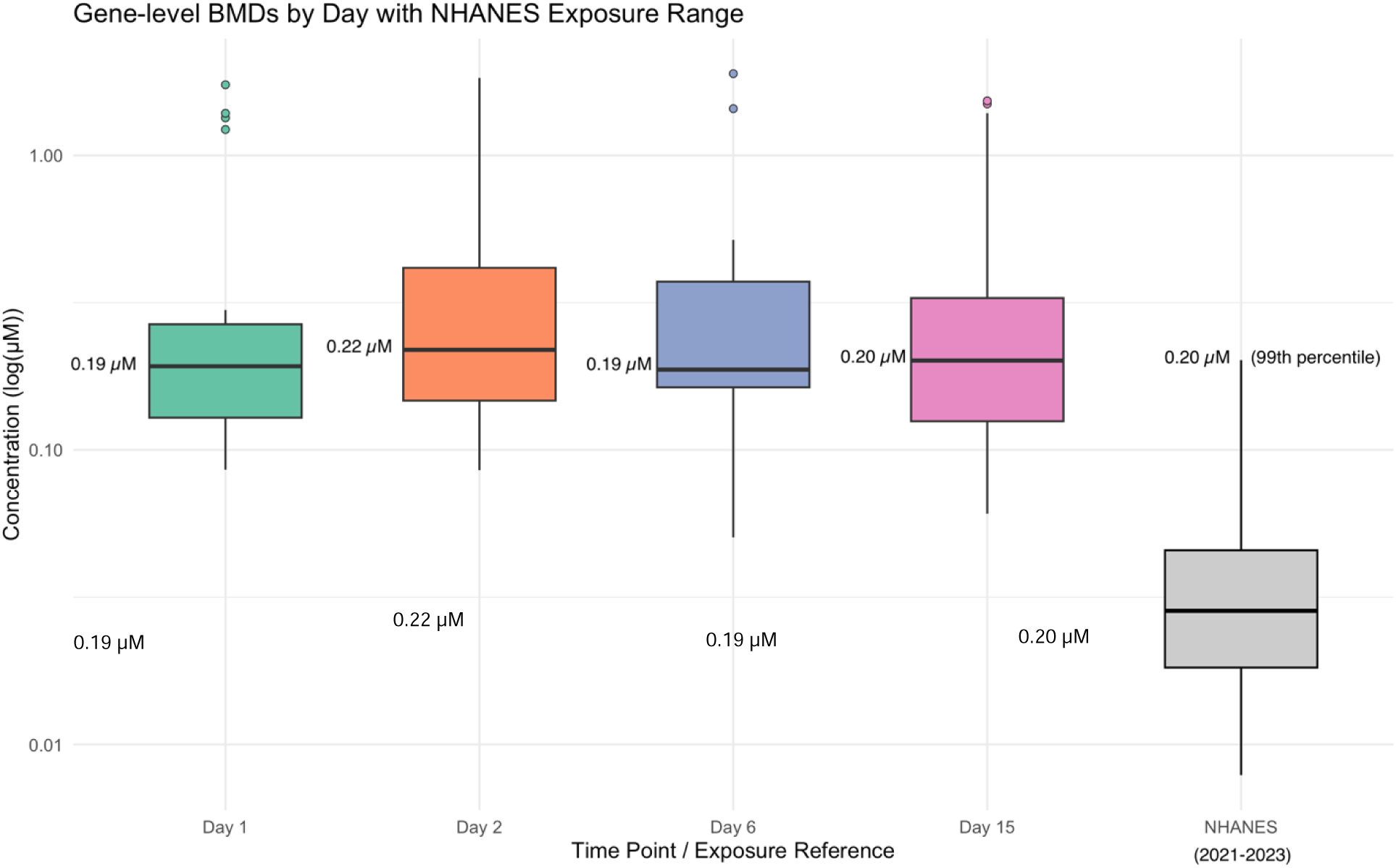
Best BMC across days of differentiation. BMCs for each day of differentiation (1, 2, 6, and 15) were chosen using the best fit model for each concentration– response relationship per gene (BMR was set to 1 standard deviation relative to control response). Each boxplot (per day) represents all genes showing a BMR. For BMC boxplots, the boxes represent the interquartile range (IQR) with median shown by the center line, and whiskers extreme to the extreme values within 1.5 x IQR. The 5^th^ boxplot shown here shows the weighted NHANES blood Pb distribution in the United States 2021-2023 as a custom weighted percentile boxplot with the box representing the IQR and the whiskers representing the 1^st^ and 99^th^ percentiles.

**Table 3:** Upregulated and downregulated BMC genes per day. Genes fitting a benchmark concentration response with directionality in response (upregulated/downregulated) derived from the “Best adverseDirection” of the winning model for a given gene.

| Day | Downregulated | Upregulated | Total |
| --- | --- | --- | --- |
| 1 | 10 | 18 | 28 |
| 2 | 23 | 25 | 48 |
| 6 | 5 | 8 | 13 |
| 15 | 14 | 17 | 31 |

In order to better understand how Pb concentration relates to the number of pathways affected, we further carried out functional classifications analysis on BMC data for the three Gene Ontology sets (Biological Process, Molecular Function, and Cellular Compartment) pathways. An accumulation plot was created to visualize the progressive increase in the number of dysregulated pathways or gene sets exhibiting BMRs across Pb concentrations for each day (**Figure 5)**. The data were filtered for significance using a Fisher’s exact two-tailed -value of < 0.05. All pathways are presented in **Table S3**, and the top 30 terms across days are shown in **Table 4**. Keywords were selected for as described above (cardiac, development, epigenetics, metabolism, and mitochondria). These selected data were plotted, showing changes only on days 1, 2, and 6, and notably showing downregulation of pathways cardiac myofibril assembly, muscle cell proliferation, and striated muscle cell proliferation for day 2 (**Figure 6**).

**Figure 5:**
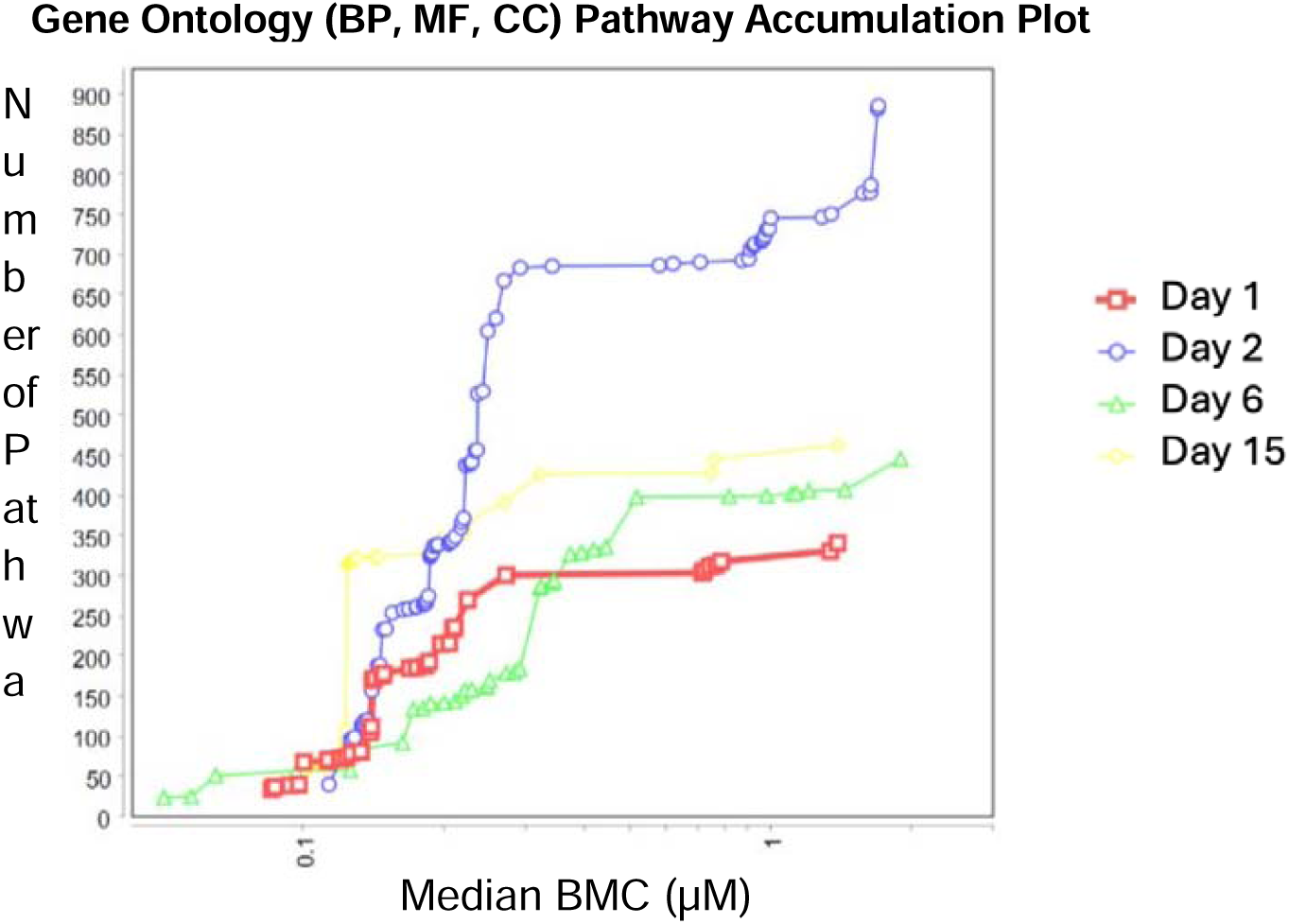
Accumulation plots for all GO (BP, MF, and CC) pathways for each day of differentiation. BP = biological processes; MF= molecular function; CC= cellular compartment. Day 1 shown in red, day 2 shown in blue, day 6 shown in green, and day 15 shown in yellow. Each plotted point represents number of pathways activated at each median BMC.

**Figure 6:**
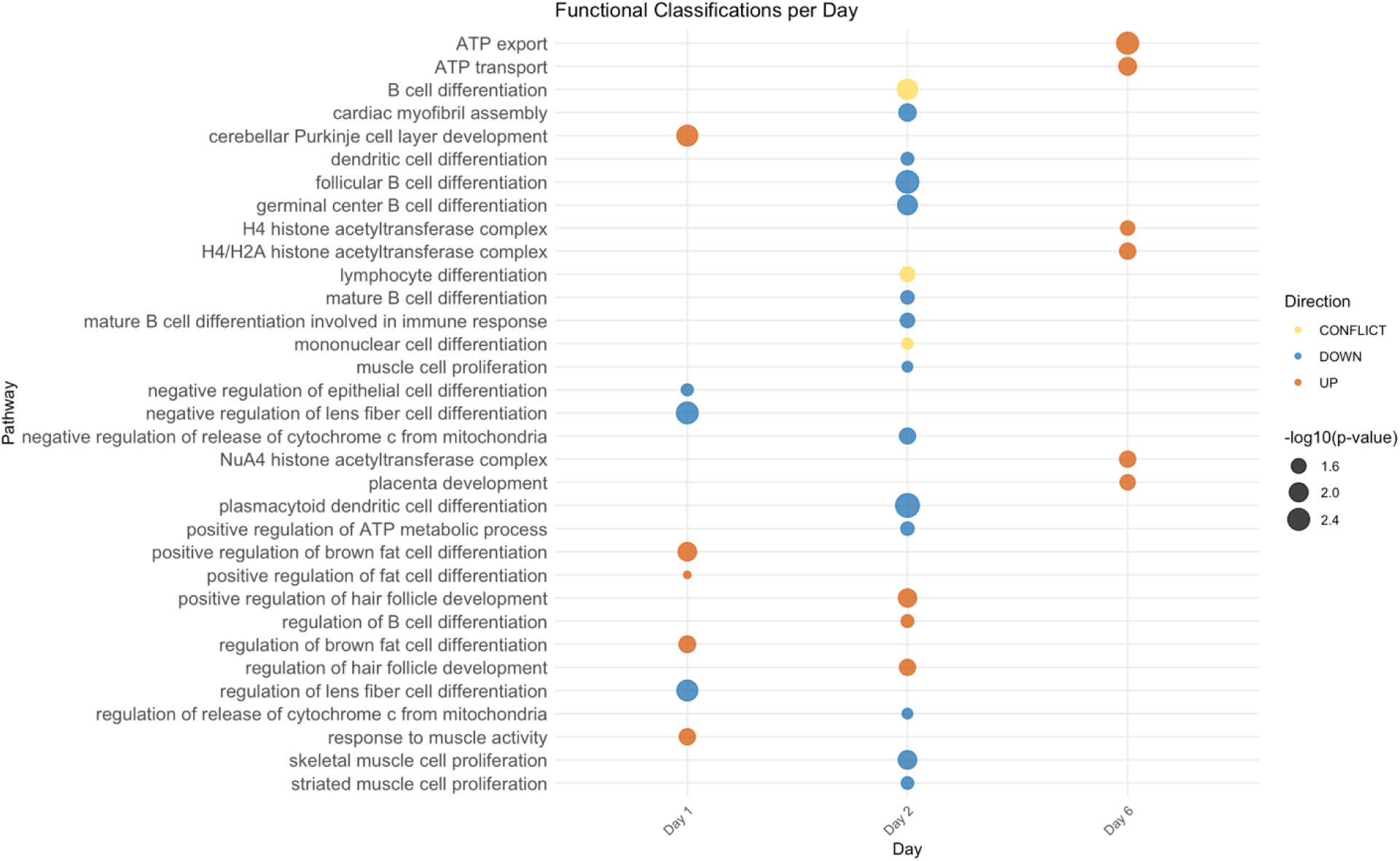
BMC modeled selected pathway changes. Using pathway data for GO pathways related to cardiac, developmental, epigenetic, metabolic, and mitochondrial responses were selected for and plotted. The size of the dot indicates relative significance (-log10(p-value)) from fisher’s exact test, with bigger circles indicating greater statistical significance. Color indicates upregulation (orange), downregulation (blue) or conflict (both upregulation and downregulation indicated by gene expression changes; yellow).

**Table 4:** Top 30 GO Terms for BMC modeled data. The top 30 pathways ranked by Fisher’s Two Tailed Exact p-value for the results for GOBP, GOCC, and GOMF for days 1, 2, 6, and 15 of ventricular differentiation, including overall direction of regulation of the effect (UP, DOWN, or CONFLICT; the latter of which means that different genes within a given pathway showed both increased and decreased expression following Pb exposure).

| **Description** | **BMC Median (μM)** | **Fisher's Exact Two Tail** | **Overall Direction** | **Day** |
| --- | --- | --- | --- | --- |
| positive regulation of cytokine production | 0.248298 | 1.06E-04 | DOWN | 2 |
| positive regulation of lymphocyte migration | 0.757676 | 1.90E-04 | CONFLICT | 15 |
| axonemal dynein complex | 1.634685 | 2.15E-04 | DOWN | 2 |
| regulation of type I interferon production | 0.248298 | 2.61E-04 | DOWN | 2 |
| regulation of lymphocyte migration | 0.757676 | 5.00E-04 | CONFLICT | 15 |
| positive regulation of interferon-beta production | 0.2064655 | 7.19E-04 | CONFLICT | 2 |
| axonemal dynein complex assembly | 1.634685 | 7.19E-04 | DOWN | 2 |
| positive regulation of mononuclear cell migration | 0.757676 | 9.13E-04 | CONFLICT | 15 |
| integrin-mediated signaling pathway | 0.745155 | 9.53E-04 | CONFLICT | 15 |
| uropod organization | 0.124272 | 9.96E-04 | UP | 15 |
| sodium:iodide symporter activity | 0.126313 | 9.96E-04 | UP | 2 |
| cellular response to Thyroid stimulating hormone | 0.126313 | 9.96E-04 | UP | 2 |
| dynein complex | 1.634685 | 0.0013085 | DOWN | 2 |
| regulation of cytokine production | 0.248298 | 0.0013331 | DOWN | 2 |
| 3'-tRNA processing endoribonuclease activity | 0.0856875 | 0.0013945 | DOWN | 1 |
| thymocyte migration | 0.124272 | 0.0014939 | UP | 15 |
| regulation of interferon-beta production | 0.2064655 | 0.0015599 | CONFLICT | 2 |
| response to type II interferon | 1.1072565 | 0.0016712 | DOWN | 6 |
| negative regulation of vesicle fusion | 0.124272 | 0.0019915 | UP | 15 |
| response to erythropoietin | 0.269185 | 0.0019915 | UP | 15 |
| cellular response to erythropoietin | 0.269185 | 0.0019915 | UP | 15 |
| plasmacytoid dendritic cell differentiation | 0.248298 | 0.0019919 | DOWN | 2 |
| iodide transmembrane transport | 0.126313 | 0.0019919 | UP | 2 |
| response to Thyroid stimulating hormone | 0.126313 | 0.0019919 | UP | 2 |
| regulation of mononuclear cell migration | 0.757676 | 0.0021547 | CONFLICT | 15 |
| DNA-binding transcription repressor activity, RNA polymerase II-specific | 0.248298 | 0.002173 | DOWN | 2 |
| DNA-binding transcription repressor activity | 0.248298 | 0.0023901 | DOWN | 2 |
| positive regulation of leukocyte migration | 0.757676 | 0.0024002 | CONFLICT | 15 |
| SNARE binding | 0.1689865 | 0.002457 | CONFLICT | 1 |
| phagolysosome assembly | 0.124272 | 0.0024888 | UP | 15 |

## Discussion

Developmental Pb exposure is associated with adverse cardiovascular health outcomes in humans, but how Pb disrupts the transcriptome during early development is largely unknown. By exposing cells to Pb during iPSC-to-cardiomyocyte differentiation, we quantified transcriptomic and concentration-responsive effects. To our knowledge, this is the first report showing that Pb exposure causes widespread disruption of developmental and stress-related transcription programs during differentiation of human stem cells into cardiomyocytes, and that these effects persist after cessation of exposure. This study paves the way for future mechanistic investigations of how Pb exposure relates to the developmental origins of cardiovascular disease.

### Exposure to Pb during differentiation alters expression of genes related to heart development and disease

Pb exposure during differentiation of iPSC into ventricular cardiomyocytes led to DEGs that were largely unique for each day of differentiation. Two genes of interest were differentially expressed with Pb exposure on both days 2 and 6: *FOXA2* and *CCKBR* (**Table S1** and **Figure S2**). *FOXA2* (Forkhead box A2 transcription factor) was found downregulated with 5 µM Pb on both days and has been shown to be required for ventricular (vs atrial) cardiac differentiation.[32] This shows that Pb may shift cell fates during differentiation and thus could have implications for cardiac structure, function, and disease risk. It also highlights the need to examine chamber specific differences in Pb cardiotoxicity in purified chamber specified cardiomyocytes, a topic currently being explored in our lab. Interestingly, *CCKBR* (the cholecystokinin B/gastrin receptor) was also downregulated on both day 2 and 6 by the 5 µM dose. Prior work has demonstrated *CCKBR* may be localized to the right atria in mice, as well as upregulated in models of myocardial hypertrophy in rats.[33, 34]

The highest number of DEGs occurred on day 6 at the 5 µM dose, with 7 genes upregulated and 70 downregulated. Of those with a |log2FC| > 1, several were related to cardiac dysfunction and disease. *FAS*, or fatty acid synthase, was upregulated with 5 µM Pb exposure on day 6. *FAS* is often upregulated in heart failure, plays a role in pathological cardiac remodeling, and has been identified as a potential therapeutic target for treating cardiac hypertrophy and heart failure.[35, 36] In the context of cardiac differentiation, day 6 corresponds to cardiac progenitor cells. Upon investigating other genes indicative of cardiac remodeling, we saw no changes in cardiac lineage markers *TBX5* nor *NKX2-5* in these cells (analyzed by qRT-PCR; data not shown). *NQO1,* NAD(P)H: quinone oxidoreductase 1, was also upregulated by Pb in these cells, and inhibits generation of reactive oxygen species and cardiovascular injury.[37] Thus, Pb upregulation of *NQO1* may reflect a compensatory response to Pb-induced oxidative stress.[38] *PLVAP*, Plasmalemma vesicle associated protein, was downregulated by Pb exposure, and it is an important endothelial factor known to play a critical role in heart development and repair, particularly in mouse models.[39] Taken together, DEGs present in day 6 cells exposed to 5 µM Pb indicate activation of regulatory mechanisms involved in cardiac development and pathology. Further mechanistic studies on how Pb dysregulates these genes during differentiation are needed in order to understand their roles and relationship with possible cardiac dysfunction.

Another interesting pattern was that of the 143 DEGs identified across all conditions, 37 did not have an assigned Human Genome Organization gene symbol and were denoted by Ensembl gene identifiers, suggesting that these loci may correspond to incompletely characterized gene features, consistent with growing evidence that the human transcriptome remains incompletely annotated and includes many poorly characterized transcribed loci. Of the 143 unique DEGs, 118 were annotated using biomaRt as protein coding, with 19 annotated as long noncoding RNA, 4 as transcribed unprocessed pseudogenes, 1 to be experimentally confirmed (TEC), and 1 as a miscellaneous RNA (**Table S4**). Further, 33 of these were either novel proteins or transcripts, 24 of which were DEGs for day 1 alone (and of these, 7 were shared between the 0.5 and 5 µM treatments; **Table S4**). Taken together, these results suggest that Pb exposure alters both well-annotated protein-coding genes as well as a substantial number of less well characterized and non-coding transcripts, further highlighting the need for additional mechanistic studies relating gene expression changes to dysregulation of cardiac development, particularly at early timepoints.

### Exposure to Pb during differentiation alters pathways related to ribosomal function and heart development

Of the top 30 pathways in the unbiased GO analysis, 18 were related to ribosomal structure or function (**Table 4**). Given the essential role ribosomes play in protein synthesis, they are critical for normal heart development and function.[40] Furthermore, ribosomal dysfunction has been implicated in various CVDs, including hypertrophy, myocardial infarctions, and atherosclerosis.[40] Thus, this data calls for additional mechanistic investigation of ribosomal alterations induced by Pb exposure, including protein studies and functional assays of ribosome activity.

Both modelling approaches (RNA-Seq pathway analysis and BMC data) indicate that pathways related to heart development may be dysregulated by Pb exposure. Dysregulated pathways included cardiac chamber morphogenesis (day 6 by 5 µM Pb), striated muscle tissue development (day 15 by 5 µM Pb), stem cell differentiation (day 6 by 5 µM Pb), heart development (day 1 by 0.5 µM Pb), cardiac chamber development (day 15 by 5 µM Pb), heart morphogenesis (day 15 by 0.5 and 5 µM Pb), cardiac chamber formation (day 6 by 0.5 µM Pb), and cardiocyte differentiation (day 6 by 0.5 µM Pb; **Figure 3**). In the BMC analysis, cardiac myofibril assembly, muscle cell proliferation, and striated muscle cell proliferation were downregulated on day 2 of differentiation (**Figure 6**). To examine cardiac differentiation efficiency, counts of *TNNT2*, *ACTN2*, and *TTN* were plotted, showing robust expression across all treatments with no significant differences, indicating Pb exposure did not affect differentiation efficiency (**Figure S4**). Overall, although differentiation into cardiomyocytes was maintained, multiple lines of evidence indicate that Pb exposure causes both early and sustained disruption of pathways related to cardiomyocyte development and maturation. This highlights the potential for Pb to produce long lasting effects on cardiac development, with the potential to increase risk of CVD later in life.

### Epigenetic pathways in iPSC-derived ventricular cardiomyocytes are regulated by Pb

Given the critical role that epigenetic mechanisms play in stem cell differentiation, and previous reports showing that Pb disrupts the epigenome in the heart, we examined whether epigenetic pathways were disrupted by Pb exposure.[19, 20, 41, 42] We found that epigenetic pathways were indeed disrupted by Pb exposure throughout differentiation (**Figure 3**). Overall, 23 epigenetic pathways were found to be dysregulated, with 11 of these on day 1, 6 pathways on day 6, and 6 pathways on day 15 for either dose of Pb (with some pathways overlapping between days and exposures; data not shown). Over half of these (12) were related to histone biology. In the BMC analysis, similar epigenetic pathways were also disrupted, with 2 histone pathways upregulated on day 6 (**Figure 6**). Importantly, altered histone biology has shown to be involved in heart failure, inflammation in CVD, and cardiac hypertrophy.[43, 44] Future work should examine whether these pathways are altered by Pb exposure in other patient iPSC lines, as well as determine whether we see changes in chromatin modifications and accessibility.

### Mitochondrial pathways are significantly altered by exposure to Pb during differentiation

We next assessed mitochondrial and metabolic pathways, because early life exposure to Pb is known to disrupt mitochondrial and metabolic function in the heart.[45, 46] Pb-induced changes in pathways related to mitochondria function were present throughout differentiation. These changes were more significant than any other set of selected pathways, with 47 mitochondria related pathways altered in total and the most significant changes on days 1, 6, and 15 (**Figure 3b**). Pb is known to cause mitochondrial damage in various tissues, leading to oxidative stress, and CVD can be induced by high levels of oxidative stress, although literature connecting Pb, oxidative stress/mitochondrial dysfunction, and the heart is limited.[47] Interestingly, in the BMC modeled data, both positive and negative regulation of the release of cytochrome c from the mitochondria were downregulated on day 2, a process important in initiating apoptosis and known to be dysregulated by Pb in rat liver, rat kidney, and human neurons (**Figure 6**).[48–50] As the heart is a highly energetically demanding organ, future work is warranted to understand Pb’s induction of mitochondrial dysfunction, oxidative stress, and apoptosis in the heart and determine the implications this may have for cardiovascular risk.

### Pb shows a similar benchmark dose on each day of differentiation despite different gene targets

BMC modeling of the RNA-seq data revealed that a low, human-relevant concentration of Pb has a significant effect on gene regulation, with median benchmark concentrations of around 0.2 µM for all 4 days examined (**Figure 4**). Similar to the LRPath analysis of sequencing results, BMC modelling showed complementary modelled dose-response gene expression changes in cardiac, developmental, epigenetic, metabolic, and mitochondrial pathways (**Figure 6**). Interestingly, day 2 showed downregulation of BMC modeled pathways pertaining to heart cell development, including cardiac myofibril assembly (driven by *MYLK3,* an essential gene in proper heart development and sarcomere formation) and striated muscle cell proliferation (driven by *HGF,* a gene that influences cardiomyocyte proliferation and differentiation in the developing heart).[51, 52]

Importantly, the CDC’s blood Pb reference value in children is 3.5 μg/dL and a BMC of 0.2 µM Pb is only slightly above this amount, equivalent to a blood Pb concentration of∼4.14 µg/dL.[53] Historical Pb exposure has been much higher, and recently during the 2014 Flint water crisis, children living in neighborhoods with the highest water Pb levels experienced a 6.6% increase in the percent of individuals with a blood Pb concentration above the then reference value of 5 µg/dL.[54] Interestingly, this median dose of 0.2 µM is comparable to the current 99^th^ percentile of population blood Pb concentrations in NHANES, suggesting that molecular disruptions identified in the BMC analysis is near the higher end of contemporary human exposure levels (**Figure 4**). This highlights the public health relevance of our transcriptomic data; significant gene alterations that disrupt important pathways during cardiomyocyte differentiation occur at human relevant and historically common exposure levels.

### Limitations and future directions

This study demonstrates widespread transcriptomic changes induced by human relevant Pb exposure during differentiation of human induced pluripotent stem cells into ventricular cardiomyocytes. Importantly, the cardiomyocytes generated here represent a fetal like state, and so conclusions that can be drawn related to the development of cardiovascular disease in adults are limited; however, by examining this early developmental stage, we can gain an understanding of potential developmental origins of cardiac malfunction and disease. Further, although pathway-level results differ between RNA-seq and BMC analyses, both datasets provide complementary evidence supporting the same biological direction, reinforcing the robustness of the transcriptomic changes observed. However, the median modeled BMCs on each day fell below the lowest tested concentration, requiring extrapolation beyond the experimental dose range and thus carrying uncertainty. Additional low concentration testing is necessary to validate the predicted low-dose effects. While this study represents a foundational understanding of how Pb during cardiomyocyte differentiation leads to transcriptomic changes, future studies should be carried out in a larger number of cell lines from both male and female donors in order to understand whether our findings are translatable the general population.

### Conclusions

We characterized transcriptomic changes induced by Pb throughout differentiation into ventricular cardiomyocytes and discovered widespread changes in expression of genes related to cardiac development and disease as well as developmental, epigenetic, and mitochondrial pathway alterations. Importantly, changes occurred at doses corresponding to human relevant exposure ranges. Overall, we found that Pb exposure during cardiomyocyte differentiation induces changes related to the development of cardiac pathology, paving the way for further mechanistic studies to interrogate the molecular underpinnings of these effects.

## Supplementary Information

Supplemental data for this work includes **Figures S1-S4** (below), the attached **Tables S1-S4**, and bm2 file from BMDExpress3 deposited here: https://github.com/kimbopossible/FemaleiPSC-CM_bmcdata

## Acknowledgements

We would like to acknowledge the DoGoodS-Pi Epigenetics Research Group and the Svoboda lab, including Isabelle Melis, Dr. Tomoko Ishikawa, and Tamara Jones for their laboratory and administrative support. We would also like to acknowledge the University of Michigan Advanced Genomics Core for RNA-Seq for conducting the sequencing for this study. This work was funded by three years of the Environmental Toxicology and Epidemiology (ETEP) training grant (T32 ES0077062) as well as by National Institutes of Health grants: R35 ES031686, P30 ES017885, R01 ES028802, and K01 ES03204812. Further funding was obtained from the Michigan Biological Research Initiative on Sex Differences in Cardiovascular Disease.

## Author Contributions

Study design was performed by Dr. Kimberley Sala-Hamrick under the guidance of Dr. Laurie Svoboda and Dr. Andre Monteiro Da Rocha. Material preparation, data collection and analysis were performed principally by Dr. Kimberley Sala-Hamrick. Anagha Tapaswi performed library preparation for RNA sequencing under the guidance of Dr. Justin Colacino. The first draft of the manuscript was written by Dr. Sala-Hamrick. All authors read and approved the final manuscript.

## Statements and Declarations

All authors declare that they have no conflicts of interest.

## Supplemental Figure

**Fig. S1:**
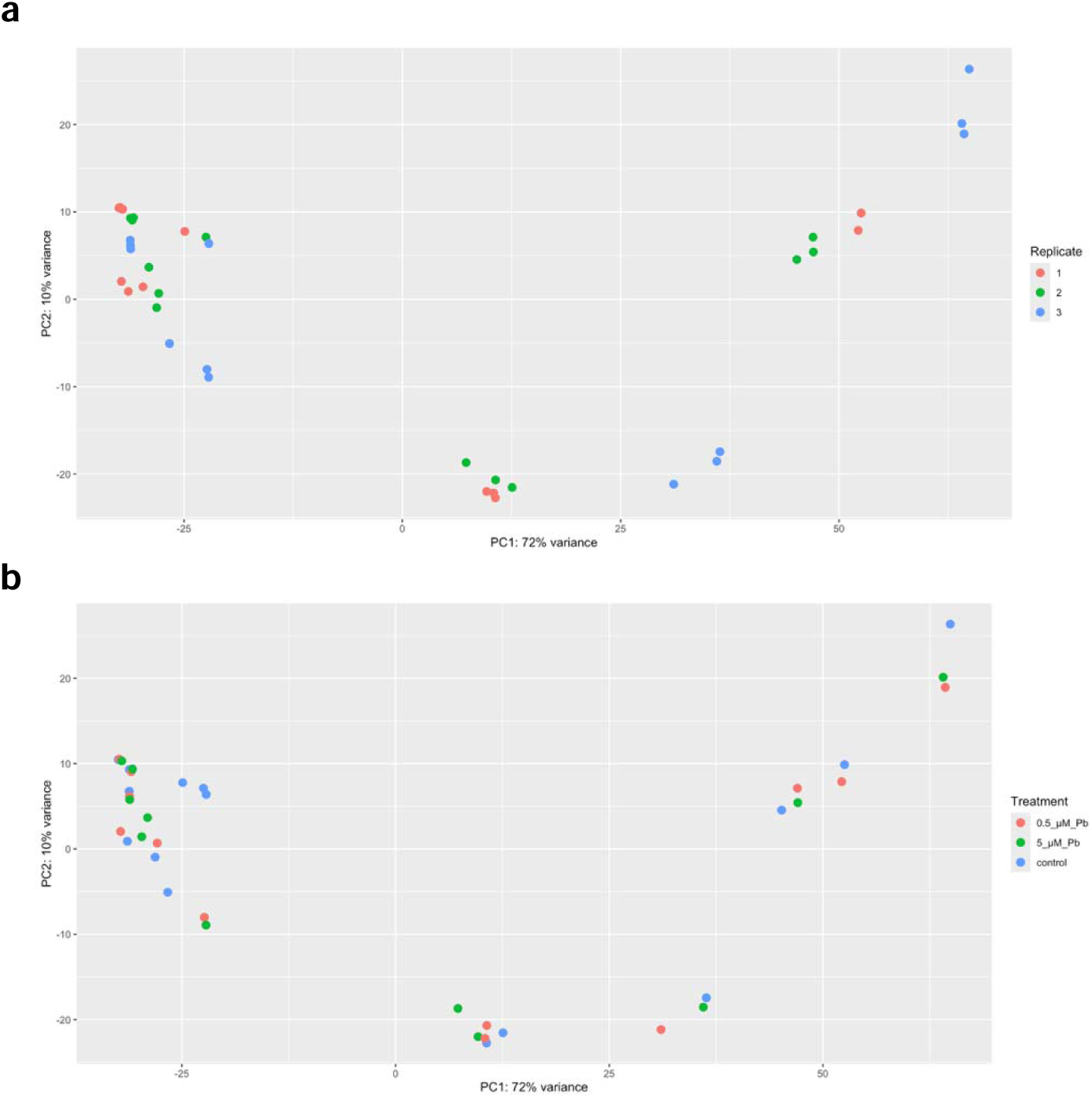
Principal component analyses for replicate and Pb treatment. PCA analyses for RNA sequencing data of female iPSCs undergoing differentiation into cardiomyocytes were plotted so each replicate (a) and each Pb treatment (b) are shown in a different color (n=3 independent experiments per treatment).

**Fig. S2:**
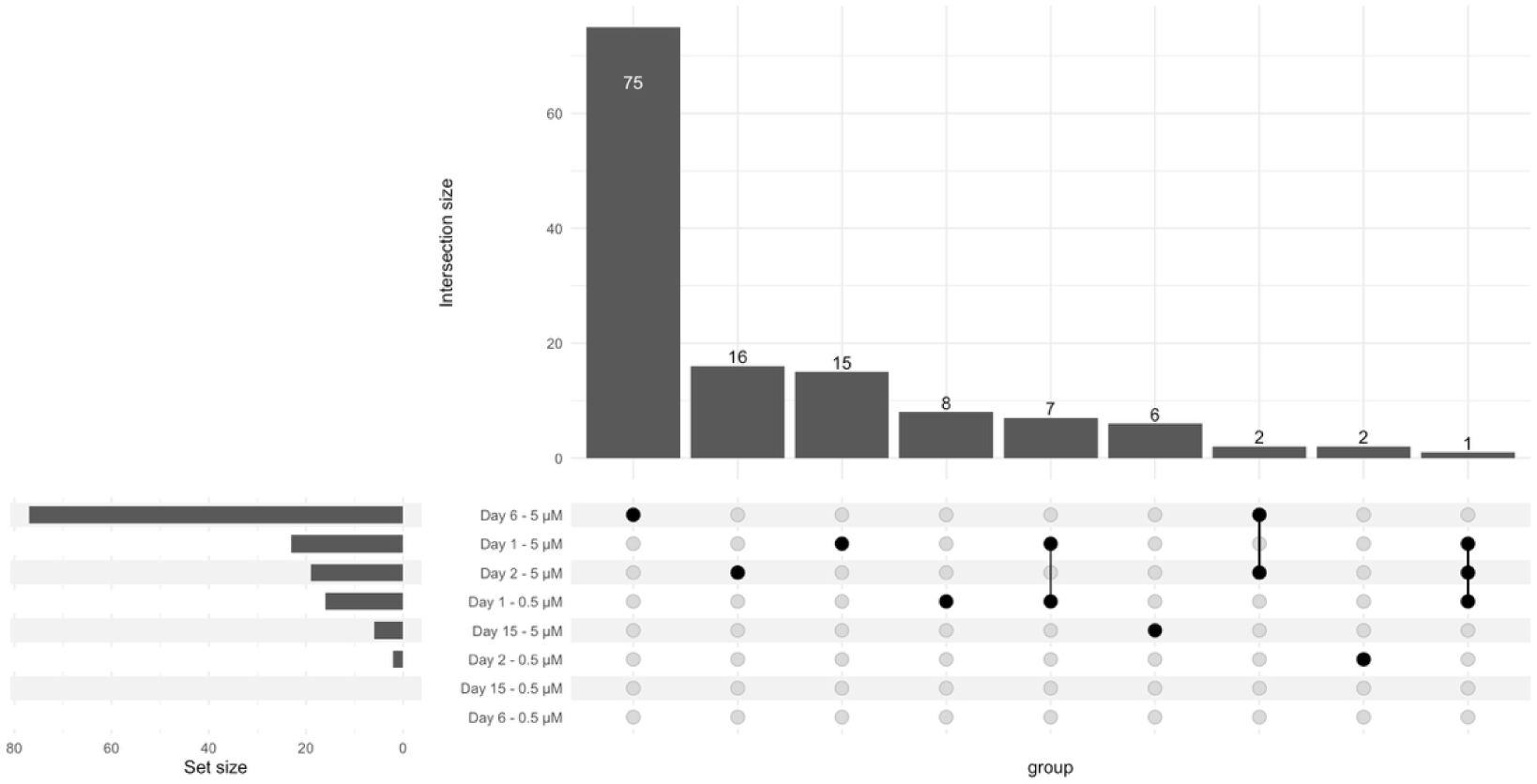
Overlaps in DEGs per day of differentiation and dose. Upset plot showing overlaps in gene expression for each day of differentiation and dose (0.5 µM and 5 µM).

**Fig. S3:**
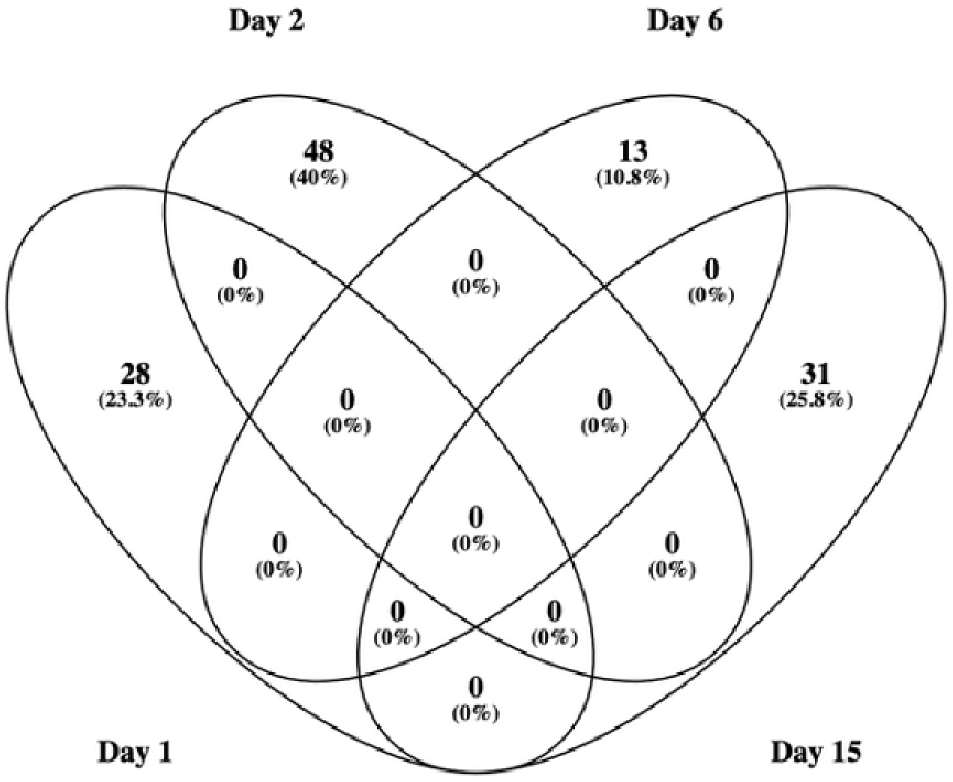
Overlap in BMC genes per day of differentiation. Venn diagram of genes with a benchmark response on days 1, 2, 6, and 15 of differentiation made by Venny 2.1.0.

**Fig. S4:**
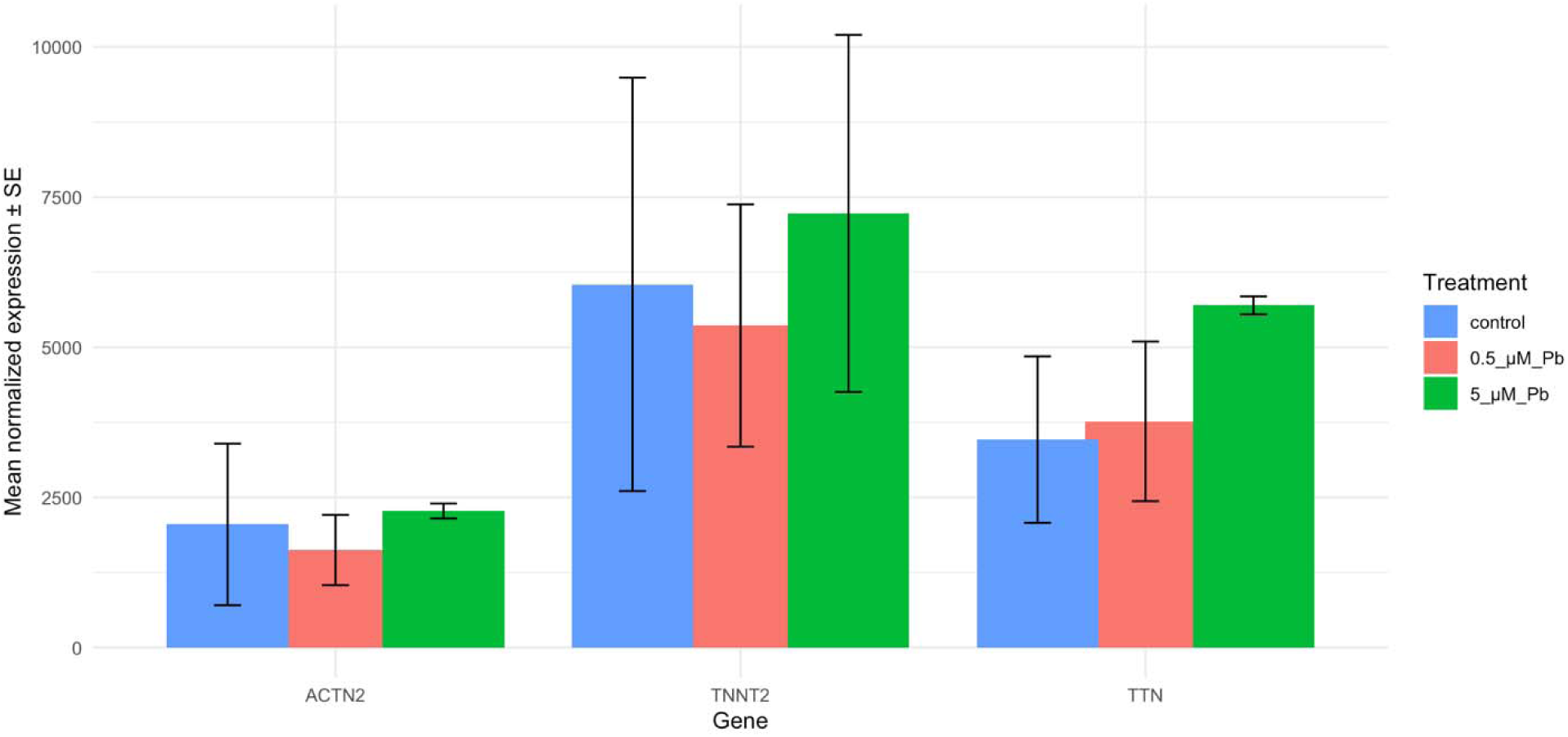
Expression of cardiac differentiation markers in day 15 cells. Mean normalized expression of *ACTN2*, *TNNT2*, and *TTN* were calculated and graphed for day 15 control and Pb exposed cells, and one-way ANOVAs found no significant differences between treatments for each gene.

